# Allosteric effects in catalytic impaired variants of the enzyme cyclophilin A may be explained by changes in nano-microsecond time scale motions

**DOI:** 10.1101/224329

**Authors:** Pattama Wapeesittipan, Antonia S. J. S. Mey, Malcolm D. Walkinshaw, Julien Michel

## Abstract

There is much debate about the mechanisms by which molecular motions influence catalysis in enzymes. This work investigates the connection between stochastic protein dynamics and function for the enzyme cyclophilin A (CypA) in wild-type (WT) form, and three variants that features several mutations that are distal from the active site. Previous biophysical studies have suggested that conformational exchange between a ‘major’ active and a ‘minor’ inactive state on millisecond timescales plays a key role in catalysis for CypA. Here this hypothesis was addressed by a variety of molecular dynamic (MD) simulation techniques. The simulations reproduce X-ray crystallography derived evidence for a shift in populations of major and minor active site conformations between the wild-type and mutant forms. Strikingly, exchange between these active site conformations occurs at a rate that is 5 to 6 orders of magnitude faster than previously proposed. Further analyses indicate that the minor active site conformation is catalytically impaired, and that decreased catalytic activity of the mutants may be explained by changes in Phe113 motions on a ns-μs time scale. Therefore previously described millisecond time scale motions may not be necessary to explain allosteric effects in CypA mutants.

---

A major goal of modern molecular biophysics is to clarify the connection between protein motions and enzymatic catalysis.^1–3^ A wide range of experimental methods, e.g. neutron scattering, X-ray crystallography, NMR, or vibrational spectroscopy have been used to characterize internalprotein motions occurring from femto second to second time scales.^4,5^ While there is broad consensus that protein motions are implicated in catalysis, there is much debate around the role of conformational changes occurring on a millisecond time scale, and several studies have linked changes in millisecond protein motions with changes in enzymatic function.^6–9^ However, it remains unclear whether such motions havea causal link to catalysis, or are merely a manifestation of the inherent flexibility of proteins over a broad range of timescales.

There have been vigorous debates about the meaning of dynamics in the context of enzymatic catalysis.^10–12^ In the framework of transition state theory, the reaction rate is given by equation 1:

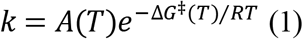

where *T* is the temperature and *R* the gas constant. The pre-exponential term *A*(*T*) includes contributions from non-statistical motions such as re-crossing or tunnelling, The exponential term involves the activation free energy of the chemical step Δ*G*^‡^(*T*). If transitions between reactant states are fast compared to the time scale of the chemical reaction, Δ*G*^‡^(*T*) is the free energy difference between the thermally equilibrated ensembles describing the reactant and transition states.^13,14^ Non-statistical motions described by *A*(*T*) have typically been found to make a small contribution to rate constants with respect to the exponential term that involves equilibrium fluctuations of the protein and solvent degrees of freedom.^15^

The current work is concerned with the connection between rates of thermally equilibrated motions, and catalysis in enzymes. Specifically, the focus is on clarifying the nature of protein motions implicated in catalysis for the well-studied enzyme cyclophilin A (CypA). CypA is a member of the cyclophilin family of peptidyl-prolyl isomerases which catalyzes the *cis*/*trans* isomerization of amide groups in proline residues.^16^ CypA plays an essential role in protein folding and regulation, gene expression, cellular signaling and the immune system. Notably, CypA is involved in the infectious activity and the viral replication of HIV-1.^17^ Accordingly, CypA has beenthe subjectof structure-based drug design efforts for decades.^18–20^ Because of its significance as a medical target, the catalytic mechanism of CypA has been the subject of extensive studies.^2,3,21–30^ Computational studies have shown that the speedup of the rate of *cis*/*trans* isomerization rate of the prolyl peptide bond is a result of preferential transition-state stabilization through selective hydrogen bonding interactions in the active site of CypA.^26,30^ Figure 1(**a**) depicts key interactions between the substrate and active site residues, whereas Figure 1(**b**) highlights the relevant *ω* angle of the substrate used to track the *cis*/*trans* isomerization reaction.

Elegant NMR relaxation experiments by Eisenmesser et al. have also characterized the existence of intrinsic motions in apo CypA that couple a ‘major’ state *M* with a ‘minor’ conformational state *m* with a rate constant *k*_*M*→*m*_= 60 s^−1^.^27^ Fraser et al. later used ambient temperature X-ray crystallographic data to determine a high-resolution structure of this CypA state *m*, revealing an inter conversion pathway with the ‘major’ state *M* that involves coupled rotations of a network of side-chains involving residues Ser99, Phe113, Met61, and Arg55. To establish there levance of this ‘minor’ state *m* to catalysis, the distal residue Ser99 was mutated to Thr99 (now only referred to as ST). Further X-ray and NMR measurements on the free enzyme confirmed that the ST mutant increased the population of the *m* state, while decreasing the conversionrate *k*_*M*→*m*_ to 1 s^−1^.^31^ Remarkably, additional NMR experiments established that this 60-fold decrease in conversion rate between *M* and *m* states in the ST mutant correlates with a ca. 70-fold decrease in bidirectional isomerization rate (*k*_*iso*_ = *k*_*cis*→*trans*_ + *k*_*trans*→*cis*_) of a model substrate with respect to wild-type (WT). The effect is comparable to rate decreases observed for mutations of key active site residues such as Arg55.^31^ More recently, two further mutants were reported in an effort to rescue the lost enzymatic activity of ST. These mutations were S99T and C115S (now only referred to as STCS), or S99T, C115S, and I97V (now only referred to as STCSIV). The two newly introduced mutants recover the enzyme activity to some extent, which correlates with an increase in *k*_*M*→*m*_ values.^32^

While this body of work suggested a link between millisecond time scale motions and catalysis in enzymes, there is currently no detailed mechanistic explanation for the decreased catalytic activity of the mutants. The present study uses a variety of extensive equilibrium and biased molecular dynamics (MD) simulations to clarify the link between catalytic activity and rates of molecular motions of CypA in wild-type and the three mutant variants.

**Figure 1.**
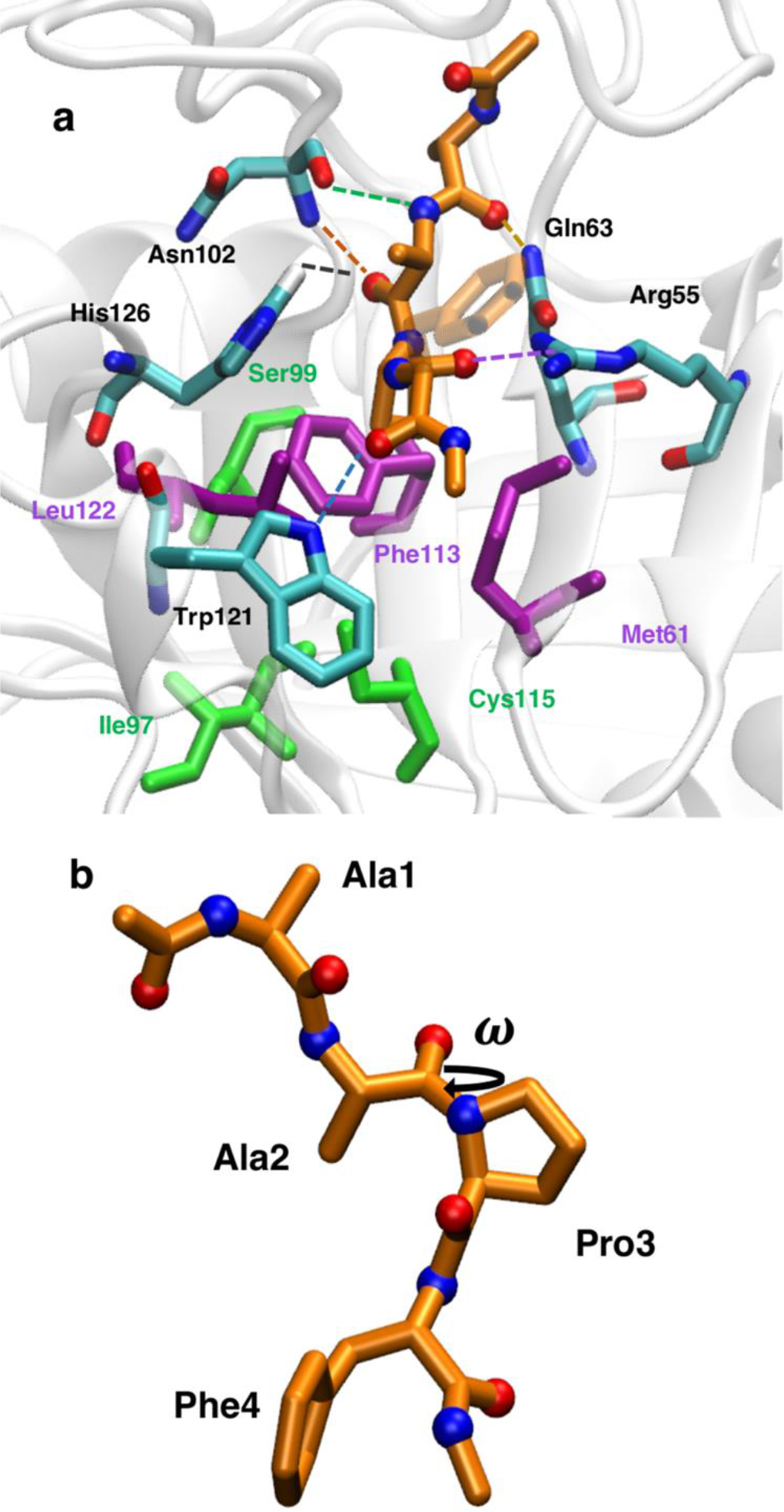
The active site of cyclophilin A (**a**) Key residues in the active site of cyclophilin A that form hydrogen bonds (cyan sticks, dashed lines) or are in contact (purple sticks) with the transition-state form of the peptide Ace-AAPF-Nme (orange sticks). For clarity, the Phe side-chain of the substrate is represented as transparent sticks. The distal residues Ser99, Cys115 and Ile97 are depicted in green. (**b**) The *ω* torsional angle of -Ala2-Pro3- is used to track the progression of the isomerization reaction between *cis* and *trans* forms.

## Results

### The proposed ‘major’ and ‘minor’ conformations exchange on timescales of nanoseconds in apo CypA

Fraser et al. have described the proposed ‘major’ and ‘minor’ states according to sets of values of *χ*_1_ (Phe113, Ser/Thr99), *χ*_2_ (Met61) and *χ*_3_ (Arg55) angles.^31,33^ The sedihedrals as well as the side-chaindihedrals *χ*_1_ of Ile97 and Cys115 were used to construct a Markov state model (MSM) to obtain quantitative information on thermodynamic and kinetic properties of the WT protein and the three experimentally studied mutants. The consistency of the MSMs was evaluated using standard protocols (See Supplementary Figures S1, S2). In the case of WT the accuracy of the MSM was additionally evaluated by back-calculation of previously reported NMR observables.^34^ The MSM yields predictions of observables that show broadly similar accuracy to that of the NMR ensembles of Chi et al. and Otter et al.(see Supplementary Figure S3).^35,36^ Thus the simulations were deemed sufficiently consistent with experimental data to warrant further analyses.

The X-ray structures of the key active site dihedrals in their dominantly populated states (if multiple occupancy is observed) are shown in Figure 2(**a**) for WT and ST, 2(**b**) for WT and STCS, and 2(**c**) for WT and STCSIV mutants. The most striking feature of the ‘major’ and ‘minor’ conformations are the rotameric states of *χ*_1_ of Phe113 from the crystal structures, which in the ‘minor’ conformation is *χ*_1_ ≈−60°. This will be referred to as the ‘*out*’ conformation. In contrast, the‘major’ state *χ*_1_ ≈60°, takes an ‘*in*’ conformation. In Figure 2, (**d**)(**e**), and (**f**) crystal structure occupancies for Phe113 *χ*_1_ are compared to the MSM-derived dihedral distributions comparing WT and ST, WT and STCS, and WT and STCSIV respectively. The simulations suggest that in apo WT the Phe113 ‘*in*’ and ‘*out*’ orientations are equally likely, which is consistent with the relatively similar occupancies of the two rotamers in the X-ray structure (occupancies = 0.63 and 0.37 respectively).^31^ In apo ST there is a significant population shift towards the ‘*out*’ orientation (*χ*_1_=−60°), and the ‘*in*’ orientation has a marginal population (ca. 1%), see Figure 2(**d**). This agrees with the X-ray structure of ST where only the Phe113 ‘*out*’ rotamer is observed (occupancy = 1.0). In the STCS and STCSIV mutants the ‘*in*’ rotamer is also destabilized with respect to wild-type but to a lesser extent (populations of ca. 16% and 17% respectively). Though only one ‘*out*’ rotamer was resolved in the X-ray structure of STCS (Figure 2(**e**)), a major ‘*out*’ and a minor distorted ‘*in*’ rotamer (*χ*_1_=+31°, occupancy 0.21) are observed in the X-ray structure of STCSIV(Figure 2(**f**)). Rotamers of other side-chain dihedrals of the key residues for all WT and mutants are found in Supplementary figures S4 and S5.

**Figure 2.**
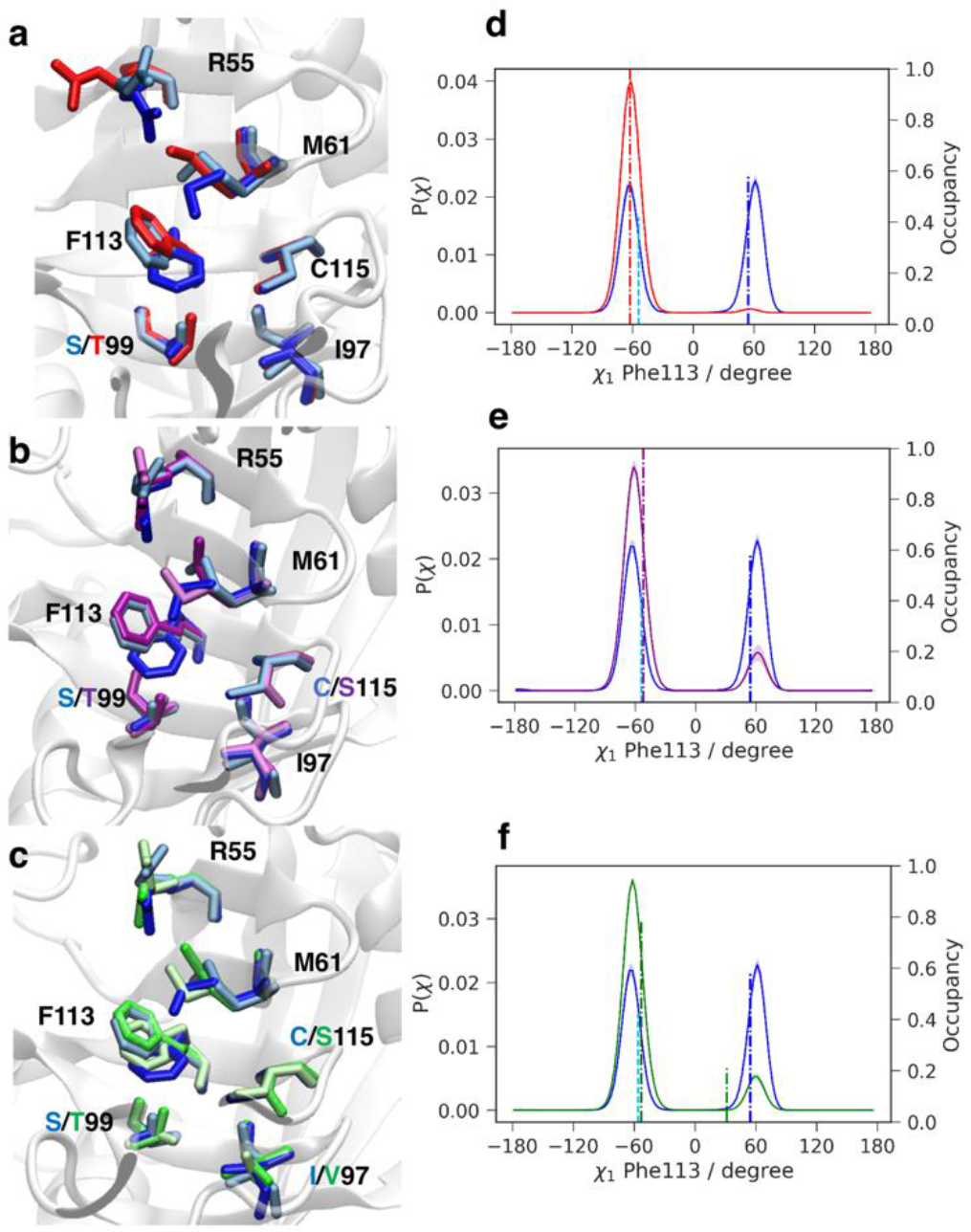
Comparison of X-ray and MSM-derived conformational preferences of Phe113. The X-ray structures of WT (blue and cyan) are compared with ST (red) (**a**), STCS (purple) (**b**) and STCSIV (dark and light green) **(c**).^31,32^ The MSM-derived probability distributions of χ_1_ in Phe113 for WT and ST (**d**), STCS (**e**) and STCSIV (**f**) are depicted as solid lines. The X-ray crystallography χ_1_ values are depicted for WT and mutants with their respective occupancies as dashed lines.^31,32^

Surprisingly the Phe113 *χ*1 dihedral was observed to flip frequently in MD trajectories of 200 ns duration (Supplementary Figure S6), suggesting faster motions than what NMR experiments have suggested. Therefore the MSM was used to obtain quantitative information on transition rates between ‘*in*’ and ‘*out*’ states as defined by the Phe113 *χ*1 rotamer. Figure 3 summarises the MSM results. WT shows the fastest overall dynamics and ST the slowest, while the STCS and STCSIV mutants show an intermediate behaviour. The exchange rates vary from 238 ± 18 μs^−1^ (ST) to 47 ± 4 μs^−1^ (STCS). Remarkably these values are five orders of magnitude faster than the exchange rates that have been determined by NMR measurements for motions involving Phe113.

**Figure 3.**
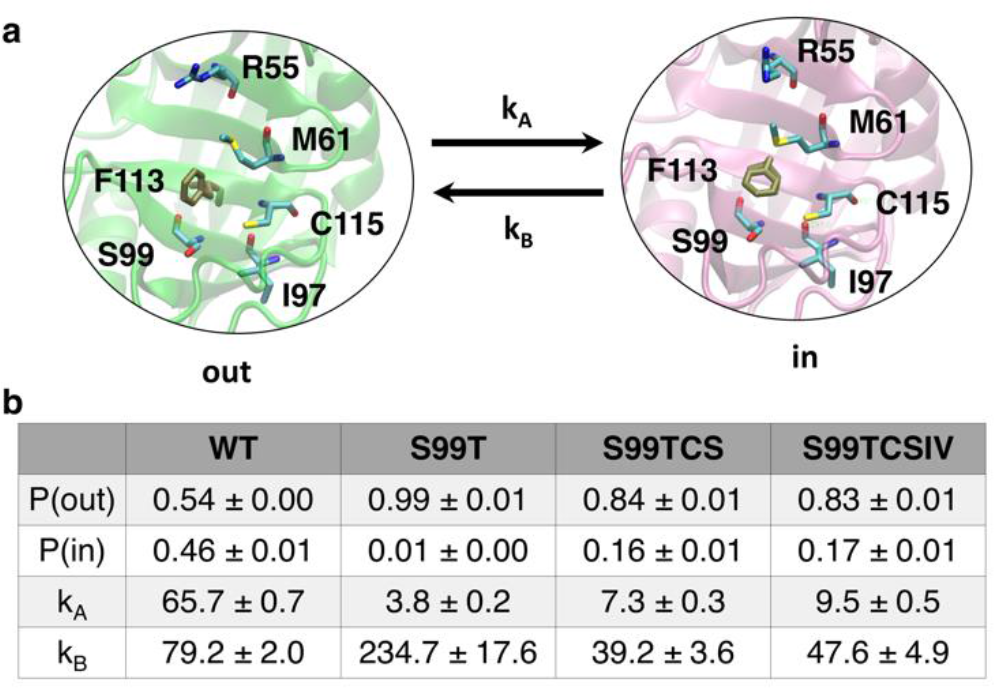
Thermodynamics and kinetics of the Phe113 in/out flip. The diagram illustrates the flip of the Phe113 *χ*_1_ dihedral between ‘*in*’ or ‘*out*’ states. Fractional populations and transition rates between ‘in’ and ‘out’ states in μs^−1^ for the four CypA variants are given in the table below the diagram. Error bars represent one standard deviation of the mean.

### No differential catalytic activity is observed in the ‘*in*’ conformations; the‘*out*’ conformation is catalytically inactive

Given that the timescales of rotations of Phe113 in the four CypA variants appear much faster than previously suggested, attention turned next to substrate bound CypA simulations. Results from umbrella sampling (US) simulations were used to quantify the isomerization free energy profile for WT and the ST mutant and investigate the role of Phe113 motions in catalysis.

The isomerization free energy profiles for WT and ST mutant with the side-chain of the Phe113 in an ‘*in*’ and ‘*out*’ conformations are shown in Figure 4(**a**) and 4(**b)** respectively. Ladani and Hamelberg have previously shown that fixed-charge classical force fields reproduce the energetics of amide bond rotation due to relatively small changes in intramolecular polarization during this process reasonably well.^28^ The calculated activation free energy for the uncatalyzed *cis*→*trans* isomerization process in water is consistent with experimental data (20.1 ± 0.1 kcal•mol^−1^ vs ca. 19.3 kcal•mol^−1^ for the related substrate Suc-AAPF-pNA at 283 K).^37,38^ The free energy profile for the substrate bound to CypA WT and ST in the ‘*in*’ conformation shows that the enzyme catalyzes the isomerization reaction in both directions via a transition state with a positive *ω* value (ca. 90-100°) equally well (Figure 4(**a**)). There is a more significant decrease in activation free energy for *trans*→*cis* (ca. −6 kcal•mol^−1^) than for *cis*→*trans* with less than 1 kcal•mol^−1^ difference between WT and ST, because the *cis* form is more tightly bound to CypA than the *trans* form. According to Figure 4(**b**), there is no catalytic benefit from the ‘*out*’ conformation of the enzyme since the activation free energy of the isomerization reaction in CypA is similar to that of the substrate in water. The calculated free energy profiles for isomerization reactions in STCS and STCSIV show a similar trend (Supplementary Figure S8).

**Figure 4.**
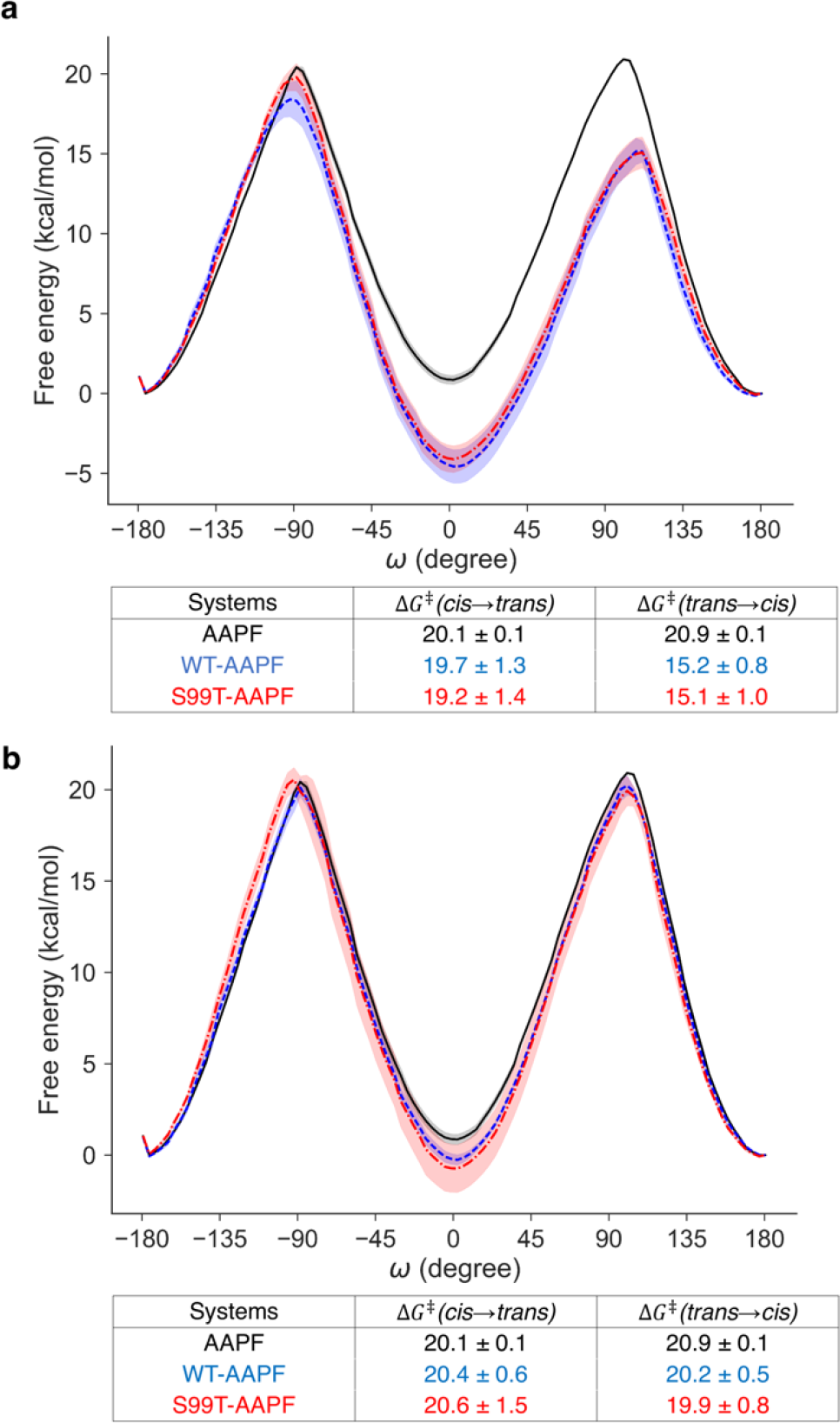
Catalytic energy profiles in the Phe113 ‘in’ and ‘out’ conformations. **(a)** Isomerization free energy profiles (in kcal.mol^−1^) for the substrate AAPF in water (black), and bound to WT (blue) or ST mutant (red) forms of CypA with starting structures in the ‘in’ conformation. The free energies of the *trans* conformation were set to zero at *ω* = 180°. Error bars represent one standard error of the mean. The table shows the activation free energies for both directions of the isomerization reaction. (**b**) same as (**a)**, but with stimulations starting in a ‘out’ configuration.

### Decreased hydrogen-bonding interactions with multiple active site residues explain transition state destabilization in the minor conformation

Further analysis of the umbrella sampling trajectories shows that for the simulations started in the ‘*in*’ configuration in both WT and ST the transition state region (*ω* ca. 90-100°) is electrostatically stabilized by more negative Coulombic interactions between substrate and binding site atoms as show in Figure 5(**a)**. Figure 5(**b**) breaks down the different contribution of active site residues, showing that Arg55, Trp121, Asn102, His126, and Gln63 are important for the stabilization of the transition state ensemble via hydrogen bonding interactions as shown in Figure 5(**e**). In contrast, Figure 5(**c**) shows that for simulations in the ‘*out*’ configuration no transition state stabilization through electrostatic interactions is observed, this is further reflected by the per-residue split of interaction energy contribution at the transition state in Figure 5(**d**) and the lack of hydrogen bond formation in Figure 5(**f**). Hydrogen bonding probabilities for simulations from the ‘*in*’ and ‘*out*’ starting conformations are shown in Supplementary Figures S9-S11. A similar picture holds for the STCS and STCSIV mutants (Supplementary Figure S12).

**Figure 5.**
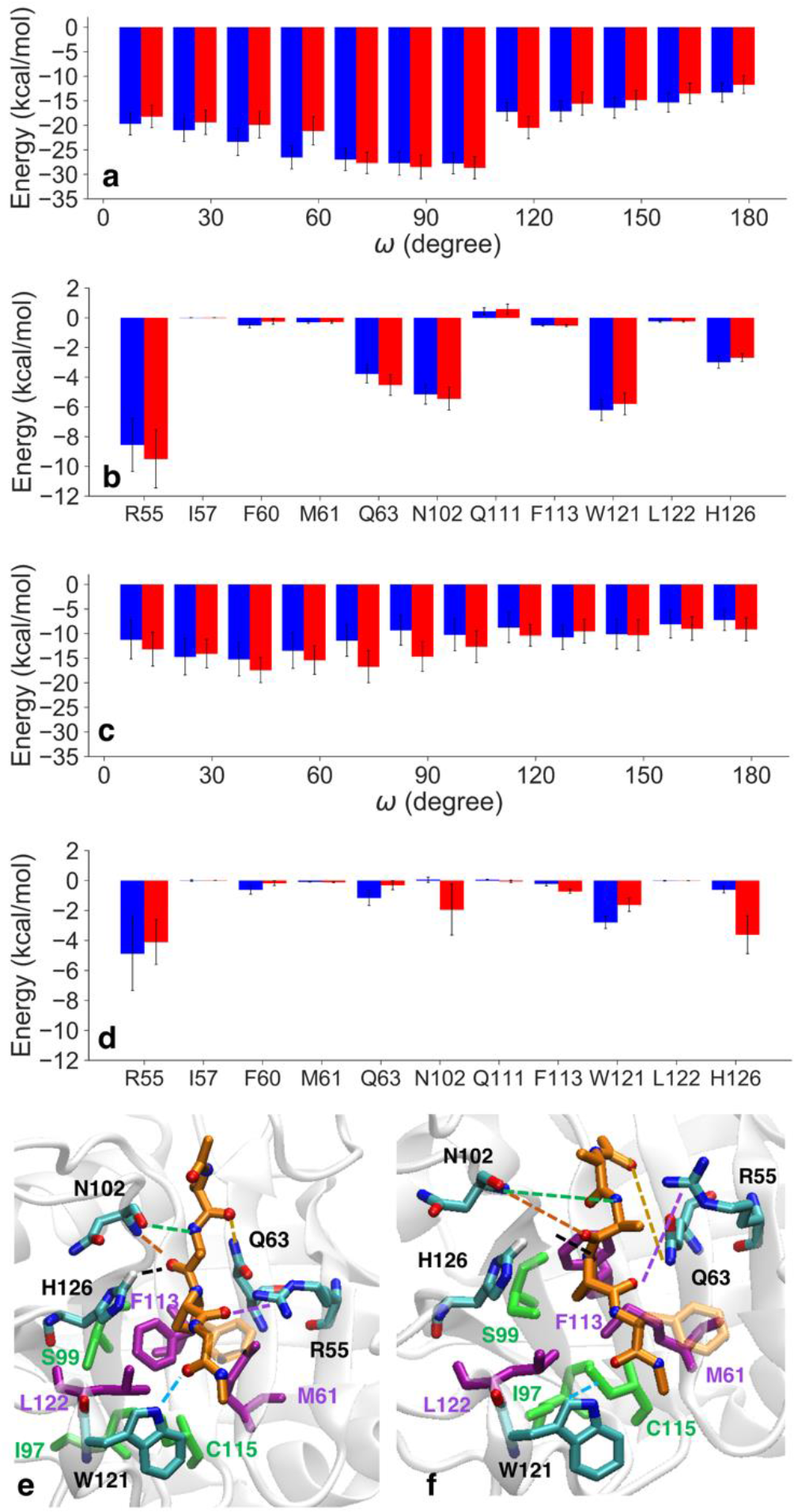
Structural basis for the differential catalytic activity of the Phe113 in and out conformations. (**a**) Electrostatic energies between the substrate and active site residues as a function of the ω angle in WT (blue) and the ST systems (red) with simulations started from the ‘*in*’ configuration. (**b**) Average electrostatic energies per-active site residues at the transition state region from (**a**). (**c**) same as (**a)** with simulations started from the ‘*out*’ configuration. (**d**) same as (**b**) with simulations started in the ‘*out*’ conformation. Error bars denote the standard error of the mean. (**e**) and (**f**) typical hydrogen bonding pattern at the transition state for simulations started in the ‘*in*’ and ‘*out*’ conformation respectively.

### Preorganization can explain observed decreases in catalytic activity of the mutants

Taken together the MSM and US data suggest a mechanistic explanation for the effect of distal mutations on the catalytic activity of cyclophilin A. In WT free form the enzyme rapidly interconverts between a catalytically active F113 ‘*in*’ form and a catalytically inactive F113 ‘*out*’ form. Because the inter conversion rate between in and out forms (ca. 7.10^7^ s^−1^) is faster than the substrate binding rate as suggested by NMR experiments (ca. 2.10^4^ s^−1^, based on *k*_on_ rate ca. 2.10^7^ s^−1^.M^−1^ and substrate concentration ca. 1 mM.)^39^ the free enzyme rapidly equilibrates between catalytically active and inactive forms before substrate binding (Figure 6(**a**)). In the mutants form, the inter conversion rates between catalytically active and inactive forms are still with in the μs^−1^ time scale, but the equilibrium is shifted towards the catalytically inactive form (Figure 6(**b**)), thus the mutants are less pre-organized than WT and the overall catalytic activity is decreased. In the case of the ST mutant and WT forms, Fraser et al. have reported bi-directional on-enzyme isomerization rates (*k*_*cis*→*trans*_ + *k*_*trans*→*cis*_)by NMR spectroscopy, and found a ratio of 68±13 between WT and ST.^31^ According to the model proposed in Figure 6 and by combining the MSM-derived populations and the US-derived activation free energies, a ratio of 12 < 46 < 176 can be derived from the simulations (see SI for details). The uncertainty from the simulations is larger than that of the measurements because small variations in activation free energies contribute large change in catalytic rates. Thus the model described in Figure 6 appears consistent with experimental data for WT and ST. No bidirectional isomerization rates have been reported for the STCS and STCSIV mutants.^32^ However, the STCS and STCSIV mutants show populations of the catalytically active Phe113 ‘*in*’ conformation that are intermediate between WT and ST, which is consistent with their increased catalytic activity with respect to ST.

A defining feature of this model is that the χ1 rotamers of a number of active-site side-chains such as Gln63, Ile/Val97, Phe113, Cys/Ser115 flip in WT and mutants on ns-μs time scales. Back-calculation of C_β_-C_γ_ order parameters shows that this effect is captured by a decrease in S^2^ values upon increasing the averaging window from 10 ns to 100 ns (Supplementary Figure S13). Motions on these timescales are too rapid to be detected by CPMG or CEST experiments that have been used extensively to study μs-ms processes in cyclophilin A.^3,25,31,40,41^ Likewise relaxation experiments cannot detect motions on this time scale as they are limited to processes occurring faster than the tumbling time τ_c_ of cyclophilin A (ca. 10 ns).^42^ Residual Dipolar Couplings (RDCs) can, however, provide information about dynamic orientation of inter-nuclear vectors on the supra-τ_c_ time scale.^43^ Such experiments have been reported for backbone and methyl-RDCs in ubiquitin.^43,44^ Therefore the model predictions can be experimentally tested with combined nuclear spin relaxation and RDC based model-free analyses coupled with a labelling scheme that resolves χ1 side-chain motions.^43,45^

**Figure 6.**
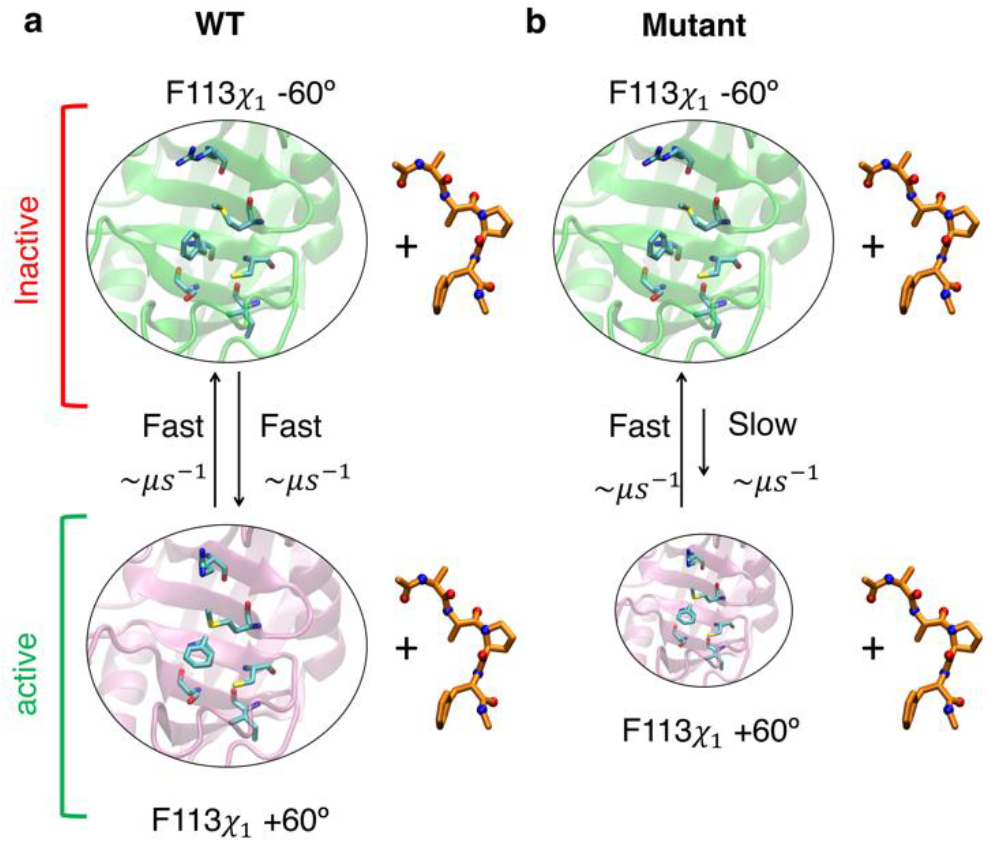
Proposed mechanism for allosteric inhibition of cyclophilin A function. **(a)** Catalysis in WT, with the favored route being the ‘*in*’ conformations, which are similarly populated to the ‘*out*’ conformations. **(b)** Catalysis in the mutants still occurs through the ‘*in*’ conformations, which have a lower population than the ‘out’ conformations.

## Discussion

This work highlights the potential of detailed molecular simulation studies to guide the interpretation of biophysical measurements for the elucidation of protein allosteric mechanisms.^46^ Previous work has suggested that exchange on millisecond timescales between conformational states in CypA are linked to its catalytic cycle,^27^ leading to a proposal for a slow exchange between a ‘major’ and a ‘minor’ state of a set of side chain rotamers linking distal residue Ser99 to active-site residues.^27,31^ The present results do not support or reject this hypothesis because the MD simulations used here do not resolve motion al processes occurring on time scales slower than microseconds. However a major finding of this study is that transitions between ‘*in*’ and ‘*out*’ rotamers of Phe 113 in WT and mutants occur on a time scale of ns-μs, thus five to six orders of magnitude faster than suggested by earlier NMR relaxation dispersion measurements.^31^ Nevertheless the simulations reproduce well the population shifts in Phe113 rotamers observed in room-temperature X-ray crystallography experiments. This suggests that the X-ray structures may have resolved motional processes occurring on a distinct timescales from the processes resolved by CPMG experiments. Indeed in reported CPMG experiments the millisecond motions of Phe113 are coupled to a large network of ca. 30 residues,^31^ whereas the χ1 rotameric flip observed in the simulations, is a largely local motion.

Nevertheless, the simulations suggest that a local ‘*in*’ to ‘*out*’ rotation of Phe113 is sufficient to abrogate catalysis in cyclophilin A, and variations of exchange parameters on the ns-μs time scale between these two conformational states appear sufficient to explain the decreased catalytic activity of the ST, STCS, STCSIV mutants with respect to WT. Therefore it is advisable to carryout additional experiments to confirm the existence of Phe113 χ1 rotations on the ns-μs time scale before causally linking catalysis to millisecond time scale motions. On the computational side, efforts should focus on advancing MD methodologies such that millisecond time scale processes observed in experiments can be resolved in atomistic details.

The contribution of protein flexibility on the ps-ns and μs-ms timescales to enzymatic catalysis has been the focus of several computational and experimental studies.^3,8,10,13,15,25,27,31,32,47^ Our work suggests that more efforts should be directed at resolving conformational processes on thens-μs time scale. This has important conceptual implications for enzyme design and optimization strategies.

## Methods

### Systems preparation

Models for apo/substrate bound human CypA of the WT and ST were prepared for MD simulations from PDB structures 3K0N (*R*=1.39 Å) and 3K0O (*R*= 1.55 Å) respectively. For apo STCS and STCSIV two structures were prepared from PDB structures 6BTA (*R*=1.5 Å) and 5WC7 (*R*=1.43Å) and also by mutating residues in WT using Schrödinger’s Maestro.^48^ For WT the major conformation of 3K0N (altloc A, occupancy 0.63) was retained. For STCS and STCSIV the residues with higher occupancy were chosen for initial structures. Supplementary Tables S1-S2 summarise all simulations conducted in this study. The proteins were solvated in a rhombic dodecahedron box of TIP3P water molecules with edges extending 1 nm away from the proteins and chloride counter-ions were added to neutralise the overall net-charge. The Charmm22* forcefield^49^ was used to describe protein atoms in the apo simulations. Steepest descent minimized was used for 50,000 steps followed by equilibration for 100 ps in an NVT ensemble, and 100 ps NPT ensemble, with heavy protein atoms restraint using a harmonic force constant of 1000 kJ·mol^−1^·nm^−2^.

Models of CypA WT and other mutants in complex with the Ace-AAPF-Nme substrate were prepared, using the amber99sb force field with optimised *ω*angle parameters for amides as reported by Doshi and co-workers was used.^50^ The crystal structure of the CypA-*cis* AAPF peptide complex(PDB ID: 1RMH)^51^ was used to obtain a suitable orientation for the substrate in the active site of WT and other mutants. PDB structure 1RMH was aligned to the structure of WT and all mutants, and the N-terminal and C-terminal ends of the proteins and substrate were capped using Schrödinger’s Maestro.^48^ In order to generate starting structures of ‘*in*’ and ‘*out*’ CypA-substrate complexes, MD simulations of CypA-substrate complexes (*cis*-conformation) were performed for 10 ns. For ST, STCS and STCSIV mutants the *χ*_1_ values of Phe113 were measured to monitor transitions between‘*out*’ and ‘*in*’ rotamers. The last snapshot structure of ‘*in*’ and ‘*out*’ complexes structures were used as input umbrella sampling calculations. For WT complexes, only the ‘*in*’ rotamer was observed in a 10-ns MD simulation. Thus, Umbrella samplings (US) simulations of *χ*_1_ (Phe113) were performed serially to generate the ‘*out*’ (*χ*_1_≈−60°) rotamer starting from the ‘*in*’ rotamer (*χ*_1_≈60°) using the software PLUMED2.^52^ Also, in order to retain the substrate in the active site, the distance between the proline ring of substrate and the phenyl rings of Phe113 and Phe60 were restrained using a force constant of 300 kJ·mol^−1^·rad^−2^. Each US simulation was performed for 5 ns. The bias parameters and the restrained variables for the US of *χ*_1_ (Phe113) are summarised in Supplementary Table S5.

### apo WT and mutant MD simulations

80 independent 200 ns MD trajectories of the apo WT, ST, STCS, and STCSIV proteins (20 each) were generated using Gromacs 5.0.^53^ For apo STCS and STCSIV the MD simulations were split between both structures prepared independently. A 2 fs time step was used, and the first 5 ns discarded for equilibration. Temperature was maintained at 300 K with a stochastic Berendsen thermostat.^54^ The Parrinello-Rahman barostat was used for pressure coupling at 1 bar.^55^ The Particle Mesh Ewald scheme was used for long-range electrostatic interactions with a Fourier grid spacing of 0.16 nm, and fourth-order cubic interpolation.^56^ Short-range van der Waals and electrostatic interaction were cutoff at 1 nm. The LINCS algorithm was used to constrain all bonds.^57^

### Markov state models

All MSM analysis was carried out with the software package pyemma version 2.3.2.^58^ The focus was on the side-chain motion of binding site residues. Details on which dihedral angles were used for TICA^59^ is given in Supplementary Table S6. A more detailed description of the Markov State model in particular with respect to best model selection is given in the SI. Clustering was done using all trajectory data from the WT and mutant trajectories using a set of 24 input coordinates, with selecting dominant coordinates usinga 90% variance in TICA for the subsequent k-means clustering. It was found that 200 clusters was necessary to separate in and out rotamers of Phe113 in ST. With the same cluster assignment for all trajectories, MSM transition matrices were estimated, using the Bayesian MSM option, and choosing lag times 0.6 ns for WT, S99T, C115S, and I97V. Means and errors of observables (e.g. populations and MFPT) were estimated from the Bayesian MSM using the provided functions in pyEMMA. Membership assignments were basedon the MSM microstate dihedral probabilities of being in the ‘*in*’ or ‘*out*’ state respectively. The microstate definition used for the MSMs is the same across the WT and all mutants. The MFPTs are estimated between the manually grouped two states depending on whether the Phe113 rotamer is ‘*in*’ or ‘*out*’ in the microstate. MSM validation and further details on the MSM can be found in the SI and in particular Supplementary Figures S1-S5.^60^

### Umbrella sampling simulations

Series of Umbrella Sampling (US) simulations ^61–63^ of the ‘*in*’ and ‘*out*’ conformers were performed to compute free energy profiles along *ω*.^26,28,64,65^ For substrate in solution, the initial structure of US was in a *trans* conformation taken from10-ns equilibration run, while all protein-substrate complexes were in a *cis* conformation. For both of ‘*in*’ and ‘*out*’ US calculations, a standard harmonic potential was used to bias the *ω* angle towards a series of target values *ω*_*k*_ spanning the interval[−180°,180°]. The force constants of the biasing potential and the spacing between *ω*_*k*_ values were adjusted by trial and error in order to obtain a good overlap between probability distributions of neighbouring *ω*_*k*_ values (Supplementary Tables S3-S4 and Figure S7). For ‘*out*’ US calculation the distances between the proline ring of substrate and the phenyl rings of Phe113 and Phe60 were restrained using a flat-bottom harmonic restraint with force constants of 200 kJ·mol^−1^·rad^−2^ and 300 kJ·mol^−1^·rad^−2^, respectively. Simulations were performed serially initially for 7 ns, with the starting conformation for a given target angle *ω*_*k*_ taken from the preceding run performed at the neighbouring *ω*_*k+Δω*_ value. Each US was then extended to 20 ns. A total of 22 (substrate in solution) or 24 (substrate bound to protein) umbrellas were used. In order to estimate uncertainties of free energy profiles six repeats of the entire procedure were performed for ‘*in*’ and ‘*out*’ US. All simulations were carried out using a PLUMED2 patched version of Gromacs 5.0 with simulation parameters identical to the previously described apo MD simulation protocols unless otherwise mentioned. The weighted histogram analysis method (WHAM) was used to produce a free energy profile from the pool of US simulations.^66^

### Other trajectory analyses

Average proton-proton distances were derived as 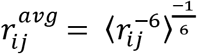 from snapshots sampled from the MSM of WT for comparison with NOEs and eNOEs-derived distance intervals.^67^ ^3^*J*(*H*^*N*^,*H*^*α*^), ^3^*J*(*H*^*N*^, *C*′), and ^3^*J*(*H*^*N*^, *C*^*β*^) were also computed using Karplus equations and backbone dihedral angle values <ϕ> and <ψ> sampled from the MSM.^68^

Interaction energies between binding site residues (Arg55, Ile57, Phe60, Met61, Gln63, Asn102, Gln111, Phe113, Trp121, Leu122 and His126) and all atoms of the substrate were analysed with the Gromacs g_energy module, using snapshots from the US simulations. The probability distribution of distances between key residues and substrate atoms during the simulations were computed using the MDAnalysis library.^69^

## ASSOCIATED CONTENT

Detailed computational protocols, supplementary figures, supplementary tables, input files for the MD calculations.

## Author Contributions

PW, ASJSM: Carried out experiments, analysed data, wrote manuscript. MDW: Analysed data, wrote manuscript. JM: Designed study, analysed data, wrote manuscript.

The manuscript was written through contributions of all authors. All authors have given approval to the final version of the manuscript.

## Data availability

All input files and scripts used for the preparation, execution and analysis of the MD, MSM and US calculations are freely available at https://github.com/michellab/CypAcatalysis_input and are also provided as Supplementary dataset S1. Other data that support the findings of this study are available from the corresponding author upon reasonable request.

## Acknowledgments

Gratitude is expressed to Fernanda Duarte for thoughtful discussions about this work.

## Graphical TOC

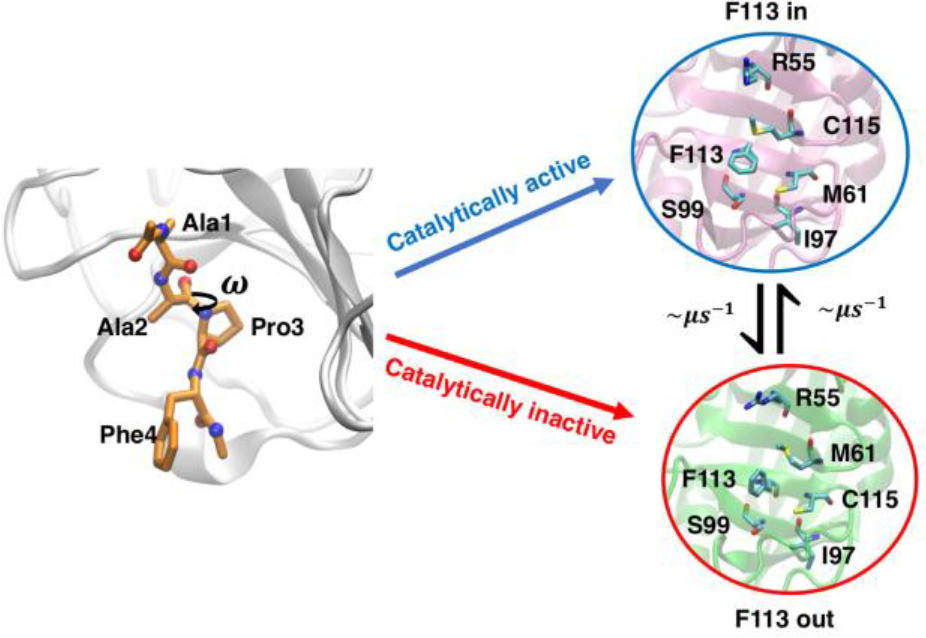

